# Root-derived *trans*-zeatin cytokinin protects *Arabidopsis* plants against photoperiod stress

**DOI:** 10.1101/2020.03.05.978221

**Authors:** Manuel Frank, Anne Cortleven, Ondrej Novak, Thomas Schmülling

**Author notes:** **Corresponding author:** Thomas Schmülling, Institute of Biology/Applied Genetics, Dahlem Centre of Plant Sciences (DCPS), Freie Universität Berlin, Albrecht-Thaer-Weg 6, D-14195 Berlin, Germany.

## Abstract

Recently, a novel type of abiotic stress caused by a prolongation of the light period - coined photoperiod stress - has been described in *Arabidopsis*. During the night after the prolongation of the light period, stress and cell death marker genes are induced. The next day, strongly stressed plants display a reduced photosynthetic efficiency and leaf cells eventually enter programmed cell death. The phytohormone cytokinin (CK) acts as a negative regulator of this photoperiod stress syndrome. In this study, we show that *Arabidopsis* wild-type plants increase the CK concentration in response to photoperiod stress. Analysis of cytokinin synthesis and transport mutants revealed that root-derived *trans*-zeatin (*t*Z)-type CKs protect against photoperiod stress. The CK signaling proteins ARABIDOPSIS HISTIDINE PHOSPHOTRANSFER PROTEIN 2 (AHP2), AHP3 and AHP5 and transcription factors ARABIDOPSIS RESPONSE REGULATOR 2 (ARR2), ARR10 and ARR12 are required for the protective activity of CK. Analysis of higher order B-type *arr* mutants suggested that a complex regulatory circuit exists in which the loss of *ARR10* or *ARR12* can rescue the *arr2* phenotype. Together the results revealed the role of root-derived CK acting in the shoot through the two-component signaling system to protect from the negative consequences of strong photoperiod stress.

## INTRODUCTION

As one of the classical plant hormones, CK regulates several developmental programs in roots and shoots (Kieber & Schaller, 2018; Werner & Schmülling, 2009) and is of crucial importance to cope with a variety of biotic and abiotic stresses (Cortleven et al., 2019).

Recently, a novel type of abiotic stress caused by a prolongation of the light period has been described and was named photoperiod stress (previously circadian stress) (Nitschke et al., 2016; Nitschke, Cortleven, & Schmülling, 2017). During a typical photoperiod stress treatment, five-weeks-old short-day (SD) grown plants were exposed to a prolonged light period (PLP). In the experimental standard setup, a PLP of 32 h was used which caused a very strong stress response, but also a PLP of 12 h (i.e. 4 h of additional light) caused a stress response (Nitschke et al., 2016). Plants exposed to photoperiod stress responded by an increased expression of numerous stress marker genes (e.g. *ZAT12* and *BAP1*) and by a decrease of genes involved in photosynthetic processes like *CHLOROPHYLL A/B BINDING PROTEIN2* (*CAB2*) about five hours after the beginning of the night following the PLP while control plants did not respond. The next day, stressed plants displayed a reduced photosynthetic efficiency and an increased percentage of water-soaked lesions that ultimately may enter programmed cell death compared to untreated and thus unaffected plants. It was found that a functional circadian clock is necessary to cope with a prolongation of the light period. Further, a particularly strong response to photoperiod stress was shown in plants with a reduced CK content or signaling suggesting that the hormone has a protective function (Nitschke et al., 2016).

Four different types of isoprenoid class CKs - *N*^*6*^-isopentenyladenine (iP), *tZ*, dihydrozeatin (DHZ) and *cis-*zeatin (cZ) - have been identified in plants and are synthesized via two different pathways requiring either adenosine mono-/di-/triphosphate (AMP/ADP/ATP) or tRNA as a precursor. Different CK metabolites can be distinguished: the bioactive free bases and the non-active ribosides, ribotides, and *O*- and *N*-glucosides (Sakakibara, 2006). In *Arabidopsis*, iP and *t*Z are the biologically most relevant CKs and are initially synthesized by the addition of dimethylallyl diphosphate (DMAPP) to AMP/ADP/ATP. This reaction is catalyzed by ADENOSINE PHOSPHATE ISOPENTENYLTRANSFERASES (IPTs) (Kakimoto, 2001; Takei, Sakakibara, & Sugiyama, 2001). Two cytochrome P450 enzymes - CYP735A1 and CYP735A2 - convert the formed iP riboside mono-/di-/triphosphate (iPRMP/iPRDP/iPRTP) molecules into *t*Z nucleotides (Takei, Yamaya, & Sakakibara, 2004). *CYP735A1* and *CYP735A2* are predominantly expressed in roots and both isoforms of the enzyme act redundantly (Kiba, Takei, Kojima, & Sakakibara, 2013). Bioactive iP and *t*Z are formed through dephosphoribosylation of iPRMP/*t*ZRMP by CK nucleoside 5’-monophosphate phosphoribohydrolase enzymes named LONELY GUY (LOGs) (Kurakawa et al., 2007; Kuroha et al., 2009; Tokunaga et al., 2012). CKs are synthesized in diverse root and shoot tissues (Miyawaki, Matsumoto-Kitano, & Kakimoto, 2004; Takei et al., 2004) and are transported through the vascular system. *t*Z-type CKs are mainly synthesized in the root and transported to the shoot via the xylem. ABCG14, an ATP-binding cassette transporter, is required for this translocation (Ko et al., 2014; Zhang et al., 2014). Root-derived *t*Z-type CKs are essential for shoot development (Kiba et al., 2013) and *t*Z and *t*ZR have distinct functions in the shoot apical meristem (SAM) and the development of leaves (Osugi et al., 2017).

Bioactive CKs activate the CK signaling cascade (Kieber & Schaller, 2014; Werner & Schmülling, 2009) by binding to ARABIDOPSIS HISTIDINE KINASE (AHK) receptors, of which *Arabidopsis* possesses three (AHK2, AHK3 and CYTOKININ RESPONSE1 (CRE1)/AHK4 (Inoue et al., 2001; Suzuki et al., 2001; Ueguchi, Sato, Kato, & Tabata, 2001; Yamada et al., 2001). Activated receptors autophosphorylate and then transfer the phosphoryl residue to AHPs (AHP1 - AHP5) (Hutchison et al., 2006). These activate type-B ARRs, which are transcription factors regulating CK-dependent gene expression (Mason, Li, Mathews, Kieber, & Schaller, 2004; Mason et al., 2005). In most cases type-B ARRs act as positive regulators of CK signaling, but one study suggested that gene regulation by type-B ARRs might be more complex (Mason et al., 2005).

The study of Nitschke et al. (2016) has shown that CK protects plants against photoperiod stress by mainly acting through the receptor AHK3 and the type-B response regulator ARR2. Further, a functional relevance of ARR10 and ARR12 as positive regulators of stress resistance was reported (Nitschke et al., 2016). However, the role of different CKs in photoperiod stress protection, the involvement of AHPs and the relationship between the different B-type ARRs has not been studied. Here, we provide evidence that plants increase their CK concentration in response to photoperiod stress and that root-derived *t*Z-type CKs protect against photoperiod stress requiring the action of AHP2, AHP3 and AHP5. The study of different type-B *arr* mutant combinations showed that ARR2, ARR10 and ARR12 together regulate the resistance to photoperiod stress.

## MATERIALS AND METHODS

### Plant material and growth conditions

The Columbia-0 (Col-0) ecotype of *Arabidopsis thaliana* was used as the wild type. The following mutant and transgenic *Arabidopsis* plants were used in this study: *abcg14-2* (Ko et al., 2014; kindly provided by Youngsook Lee); *cyp735a1-2 cyp735a2-2* (*cypDM*; Kiba et al., 2013; kindly provided by Hitoshi Sakakibara); *ahp2-1 ahp3 ahp5-2* and respective double mutants (Hutchison et al., 2006); *arr2* (GK-269G01; Nitschke et al., 2016); *arr10-5 arr12-1* and the respective *arr10-5* and *arr12-1* single mutants (Argyros et al., 2008; Mason et al., 2005). If not mentioned otherwise, seeds were obtained from The European Arabidopsis Stock Centre (NASC; http://arabidopsis.info/). The *arr2 arr10-5 arr12-1*, *arr2 arr10-5*, *arr2 arr12-1* mutants were generated by genetic crossing and the genotypes were confirmed by PCR analysis. *Arabidopsis* plants were grown on soil in a growth chamber under SD conditions (8 h light/16 h dark) as described in Nitschke et al. (2016). For photoperiod stress treatment, plants were exposed to a light period of 32 h. For CK treatment, plants were watered daily from below (ca. 150 mL/tray corresponding to ca. 4 mL/plant) with either 10 μM *t*Z (dissolved in 0.01 % DMSO), 10 μM *t*ZR (dissolved in 0.01 % DMSO) or 0.01 % DMSO (control) dissolved in water.

### Quantification of lesions

Water-soaked lesions were quantified three to four hours after the night following PLP treatment. First, the total number of fully expanded leaves (except for leaf 1 and 2 as well as cotyledons) of a plant was counted. Afterwards, the total number of limp leaves was determined (0 = no water-soaked lesion, 0.5 = less than 50 % of leaf surface water-soaked, 1 = more than 50 % of leaf surface water-soaked) and the percentage was calculated for each plant by dividing the number of limp leaves by the total number of fully expanded leaves.

### Chlorophyll fluorometry

As a measure of the response to photoperiod stress the photosystem II maximum quantum efficiency (F_v_/F_m_ ratio; Baker, 2008) was determined six to seven hours after the night following the PLP. First, healthy and lesioned leaves of several plants (three leaves per plant) were detached in a ratio reflecting the determined lesion percentage of the respective genotype in the same experiment. Detached leaves were placed in Petri dishes filled with water with the abaxial part of the leaf directly facing the water. After 20 min of incubation in darkness, pulse-amplitude-modulated (PAM) measurements were performed with the chlorophyll fluorometer FluorCam (Photon Systems Instruments). The minimum fluorescence emission signal F_0_ was recorded first and then the maximum fluorescence yield F_m_ (induced by a saturating light pulse of 1500 μmol m^−2^ s^−1^).

### RNA isolation and quantitative RT-PCR

Ca. 100 mg of leaf material was harvested into 2 mL Eppendorf tubes and shock-frozen in liquid nitrogen under white light (0 h time point) or green safety light (7.5, 15 h time points). RNA isolation was performed as described by Sokolovsky et al. (1990) with a few alterations. Briefly, frozen samples (100 mg fresh weight) were ground using a Retsch mill in pre-cooled adapters. Afterwards, samples were solved in 750 μL extraction buffer (0.6 M NaCl, 10 mM EDTA, 4 % (w/v) SDS, 100 mM Tris/HCl pH 8) and 750 μL phenol/chloroform/isoamyl alcohol (PCI; 25:24:1) solution was added. Samples were vortexed, shaken for 20 min at room temperature and centrifuged at 19.000 *g* for 5 min at 4 °C. The supernatants were transferred into fresh 1.5 mL Eppendorf tubes and CI solution was added in a 1:1 ratio. Samples were vortexed briefly and centrifuged at 19.000 *g* for 5 min at 4 °C.

Supernatants were transferred into fresh tubes and RNA was precipitated for 2 h on ice by adding 0.75 volumes of 8 M LiCl. After centrifugation at 19.000 *g* for 15 min at 4 °C, supernatants were removed and resolved in 300 μL RNase-free water. RNA was precipitated again by the addition of 30 μL 3 M sodium acetate and 750 μL absolute ethanol and incubation at −70 °C for 30 min. Samples were centrifuged at 19.000 *g* for 10 min at 4 °C and the supernatant was discarded. Pellets were washed with 200 μL 70 % ethanol and after centrifugation, pellets were dried at room temperature and resolved in 40 μL RNase-free water.

cDNA synthesis and qRT-PCR analysis were performed as described in Cortleven et al. (2016) using 500 ng of total RNA and a CFX96™ Real-Time Touch System (Bio-Rad Laboratories GmbH; Feldkirchen, Germany). All primers used in this study can be found in Supplemental Table 1 of Nitschke et al. (2016). Gene expression data were normalized against reference genes according to Vandesompele et al., 2002. *PROTEIN PHOSPHATASE2A SUBUNIT A2* (*PP2AA2*, AT3G25800), *UBIQUITIN-CONJUGATING ENZYME10* (*UBC10*, AT5G53300) and *METACASPASE 2D* (*MCP2D*, AT1G79340) served as reference genes.

### Determination of CK concentrations

For CK measurements, 100 mg fresh weight of leaf tissue per sample was collected and shock-frozen in liquid nitrogen under white light (time points during light exposure) or green safety light (time points during night). CK quantification was performed according to the method described by Svačinová et al. (2012), including modifications described by Antoniadi et al. (2015). Using 15 mg per technical or biological replicate, samples were homogenized and extracted in 1 ml of modified Bieleski buffer (60% MeOH, 10% HCOOH and 30% H_2_O) together with a cocktail of stable isotope-labeled internal standards (0.25 pmol of CK bases, ribosides, *N*-glucosides, and 0.5 pmol of CK *O*-glucosides, nucleotides per sample added). The extracts were applied onto an Oasis MCX column (30 mg/1 ml, Waters), eluted by two-step elution using 1 ml of 0.35M NH_4_OH aqueous solution and 2 ml of 0.35M NH_4_OH in 60% (v/v) MeOH solution and then evaporated to dryness *in vacuo*. CK analysis was carried out using ultra-high performance liquid chromatography-electrospray tandem mass spectrometry using stable isotope-labelled internal standards as a reference. All samples were measured in quintuplicate for each genotype and each time point.

### Statistical analysis

For CK measurements, the significance of differences between control and PLP samples was calculated with a paired Student’s t-test in Microsoft Excel^®^. For statistical analysis of all other data SAS^®^Studio (https://odamid.oda.sas.com/SASStudio) was used. Homogeneity and homoscedasticity were tested by Shapiro-Wilk and Levene tests (p ≥ 0.01) before ANOVA testing was performed followed by Tukey post hoc test. If assumptions were not met, transformations (log_2_, log_10_, sqrt, n^0.1^, n^0.4^, n^1.5^, n^7^, n^25^) were performed. Paired Wilcoxon test was performed if assumptions were still not met after transformation.

## RESULTS

### Photoperiod stress increases the CK content in wild-type plants

Plants impaired in CK biosynthesis or signaling are sensitive to photoperiod stress (Nitschke et al., 2016). To investigate whether photoperiod stress influences the CK concentration, we have measured CK in leaves of SD-grown wild-type plants exposed to a PLP of 32 h, which is the standard stress treatment used in this study (Fig. 1A). The altered light regime caused an elevated total CK concentration at the end of the PLP and in the middle of the following night (Fig 1B; time points 2 and 3). The concentration of CK free bases was elevated up to three-fold in PLP plants compared to control plants at the end of the PLP and in the middle and at the end of the following night (Fig 1C; time points 2, 3 and 4). A similar pattern was observed for the concentration of CK ribosides (Fig. 1D). CK nucleotides were increased two-fold in PLP plants compared to control plants after 16 h of additional light (Fig.1E; time point 1) and stayed elevated in PLP plants in comparison to control plants until the end of the night following the PLP (Fig. 1E.; time points 2 and 3). Concentrations of CK *O*-glucosides were elevated in PLP plants during and at the end of the night following the PLP while concentrations of *N*-glucosides did not differ between stressed and control plants (Fig. 1F, G). The increase in the sum of free bases, nucleosides and nucleotides was reflected by the increased concentrations of the respective individual iP-, *t*Z- and DHZ-type CK metabolites already during the PLP (Table S1; time points 1 and 2). In contrast, the concentrations of *c*ZR and *c*ZRMP levels were decreased in PLP plants at early time points but strongly increased at the end of the night following the PLP and the day after (Table S1; time points 1, 2, 4, 5). Taken together, photoperiod stress treatment led to an increase of all types of isoprenoid class CKs including the bioactive CKs iP and *t*Z as well as their transport forms and precursors.

**Figure 1.**
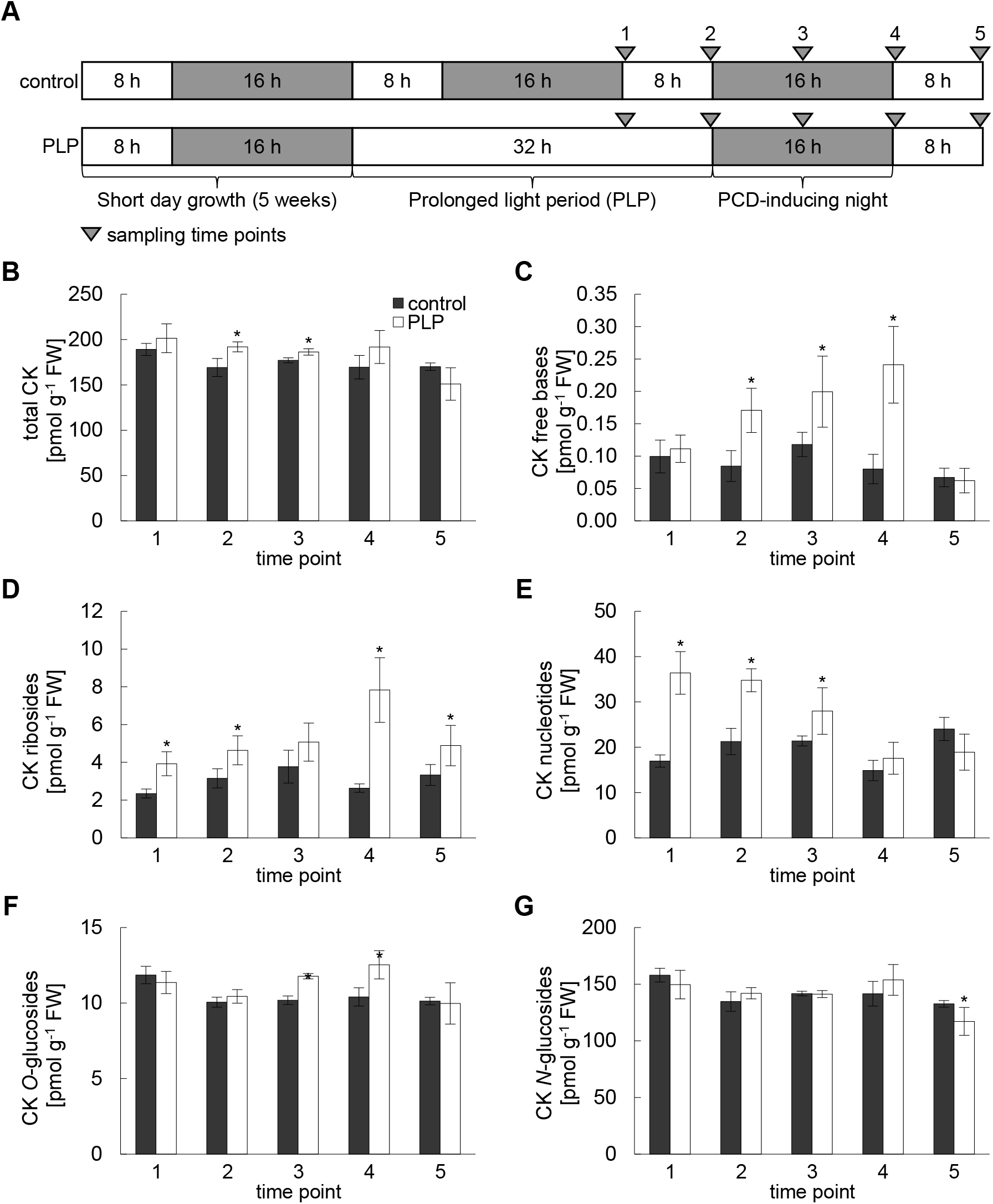
Photoperiod stress increases the CK concentration in wild-type plants. (A) Schematic overview of sampling time points for CK measurements. 5-weeks-old wild-type plants were cultivated under SD conditions and were further cultivated under these conditions (control) or were exposed to a prolonged light period (PLP) of 32 h. (B - G) Concentration of total CK (B), CK free bases (C), CK ribosides (D), CK nucleotides (E), CK *O*-glucosides (F) and CK *N*-glucosides (G) in control and PLP samples at the time points depicted in A. Stars indicate a statistically significant difference between PLP and the respective control samples at the given time point (1 to 5) in a paired Student’s t-test (p ≤ 0.05). Values are given as pmol g^−1^ FW ± SD (n = 5). The complete data set is shown in Table S1.

### Root-derived *t*Z-type CKs protect plants from photoperiod stress

Since stressed wild-type plants increased the concentration of the functionally most relevant CKs - iP and *t*Z - we wondered which of these two CKs might be protective against photoperiod stress. Therefore, we investigated the involvement of *t*Z-type CKs by exposing mutants impaired in either the biosynthesis of *t*Z-type CKs (*cypDM*; Kiba et al., 2013) or their transport from the root to the shoot (*abcg14*; Ko et al., 2014; Zhang et al., 2014) to photoperiod stress. Only results of PLP-treated plants are shown since control plants do not show any differences in Fv/Fm, lesion formation nor an altered gene expression during the course of the experimental treatment (Nitschke et al., 2016).

Over 80 % of the leaves of *cypDM* and *abcg14* mutants showed lesion formation after photoperiod stress treatment, which was a four-fold increase compared to wild-type plants (Fig. 2B, Suppl. Fig. 1A). Furthermore, photoperiod stress caused a drop in Fv/Fm to 0.35 in these mutants while wild-type leaves had an Fv/Fm value of 0.8 (Fig. 2C). The transcript abundance of the stress marker genes *BAP1* and *ZAT12* was increased in the response to stress two- to three-fold higher in the mutants as compared to wild type (Fig. 2D, E). The abundance of *CAB2* transcript was strongly decreased in all genotypes but much stronger in both mutants compared to wild type 15 h after the PLP (Fig. 2F). Summing up, these results support a protective function of root-derived *t*Z-type CKs against photoperiod stress.

**Figure 2.**
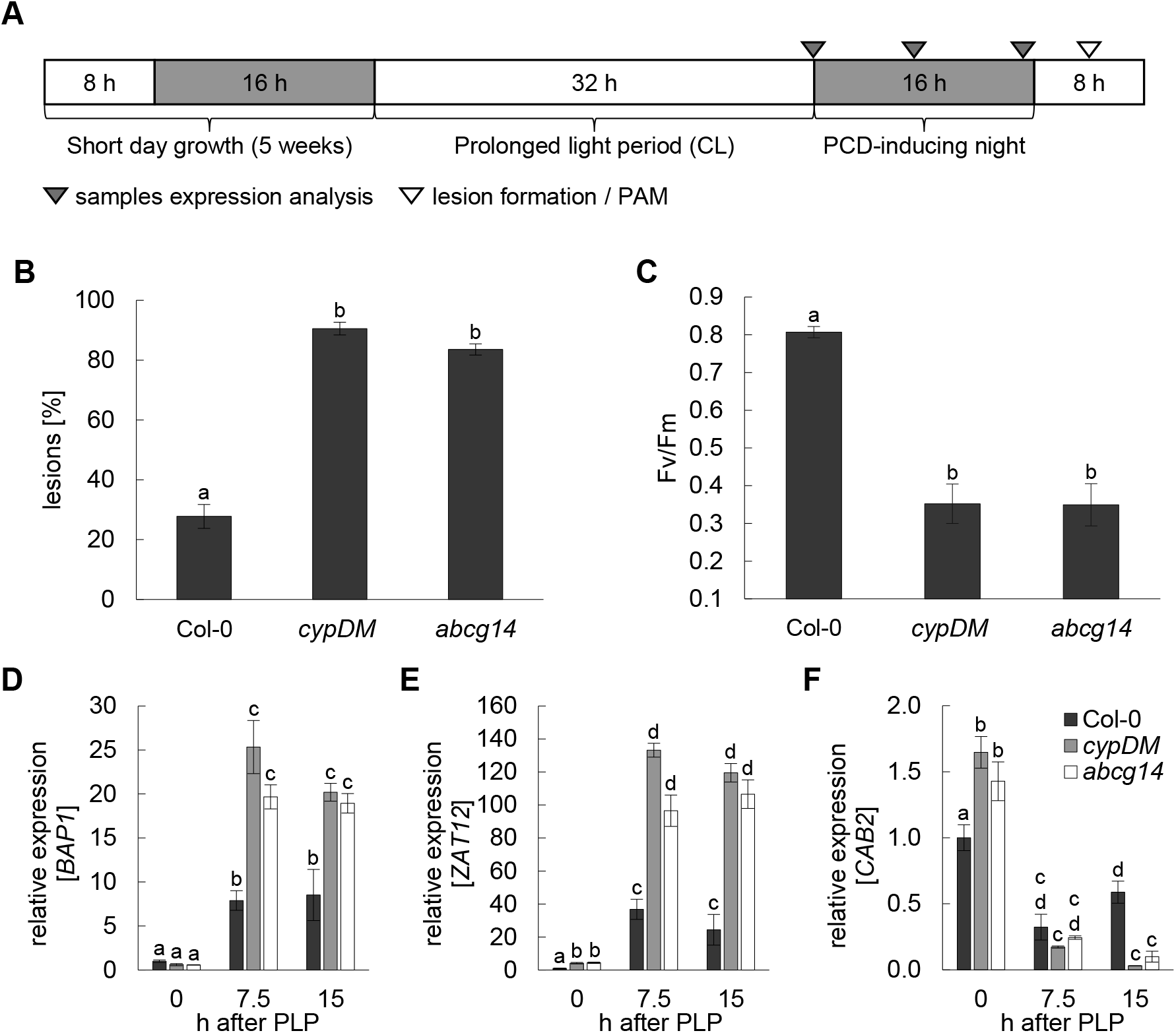
Plants deficient in *t*Z-type CKs are strongly affected by photoperiod stress. (A) Schematic overview of photoperiod stress treatment. Arrow points indicate sampling time points for the different analysis. (B) Lesion formation of leaves in 5-weeks-old Col-0, *cypDM* and *abcg14* plants the day after the PCD-inducing night (one-way ANOVA; p ≤ 0.05; n = 15). (C) Photosystem II maximum quantum efficiency (Fv/Fm) of leaves the day after the PCD-inducing night (Paired Wilcoxon test; p ≤ 0.05; n = 15). (D - F) Expression of marker genes (*BAP1, ZAT12*, *CAB2*) 0 h, 7.5 h and 15 h after PLP treatment. Letters indicate statistical groups (two-way ANOVA; p ≤ 0.05; p ≤ 0.05; n ≥ 3). The expression level of wild type at timepoint 0 h was set to 1. Error bars indicate SE. Pictures of representative plants exposed to a 24-h prolongation of the light period are shown in Fig. S1A.

### Watering of cypDM plants with *t*Z or *t*ZR reduces the response to photoperiod stress

A recent study by Osugi et al. (2017) demonstrated that under long-day conditions root-derived *t*Z has distinct functions in the shoot as compared to root-derived *t*ZR, for example in regulating the size of leaves and of the SAM. In order to dissect the role of root-derived *t*Z and *t*ZR in photoperiod stress, we watered *cypDM* plants with either 10 μM *t*Z or 10 μM *t*ZR daily during the whole cultivation period and exposed them subsequently to photoperiod stress. The effectiveness of the treatment was tested by determining the expression of CK response genes *ARR5* and *ARR6* (Supplemental Fig. S2). Expression of both genes was lower in control *cypDM* plants compared to wild type but could be rescued by application of *t*ZR and *t*Z.

Moreover, *t*ZR application reduced lesion formation in *cypDM* plants in response to photoperiod stress by about 15 % compared to untreated *cypDM* plants (Fig. 3A, Suppl. Fig. 1B). Also, the decrease in photosynthetic capacity of *t*ZR-treated plants was lower compared to untreated *cypDM* controls and almost like wild type (Fig. 3B). These results indicate that *t*ZR applied through roots has a protective effect against photoperiod stress. Watering plants with *t*Z suppressed the photoperiod stress syndrome in *cypDM* plants almost completely suggesting that also root-derived *t*Z protects plants during photoperiod stress (Fig. 3A, B). At the molecular level, DMSO treatment itself lowered the expression of stress marker genes *ZAT12* and *BAP1* (Fig. 3C, D). *t*ZR and *t*Z supplementation reduced the induction of these genes as well. The rescue of gene regulation as a response to photoperiod stress by *t*Z was particularly evident in the case of *CAB2* (Fig. 3E). In summary, supplementation experiments indicated that lesion formation, the decrease in photosynthetic capacity and the transcriptional response can be rescued to a different extent by *t*Z and *t*ZR.

**Figure 3.**
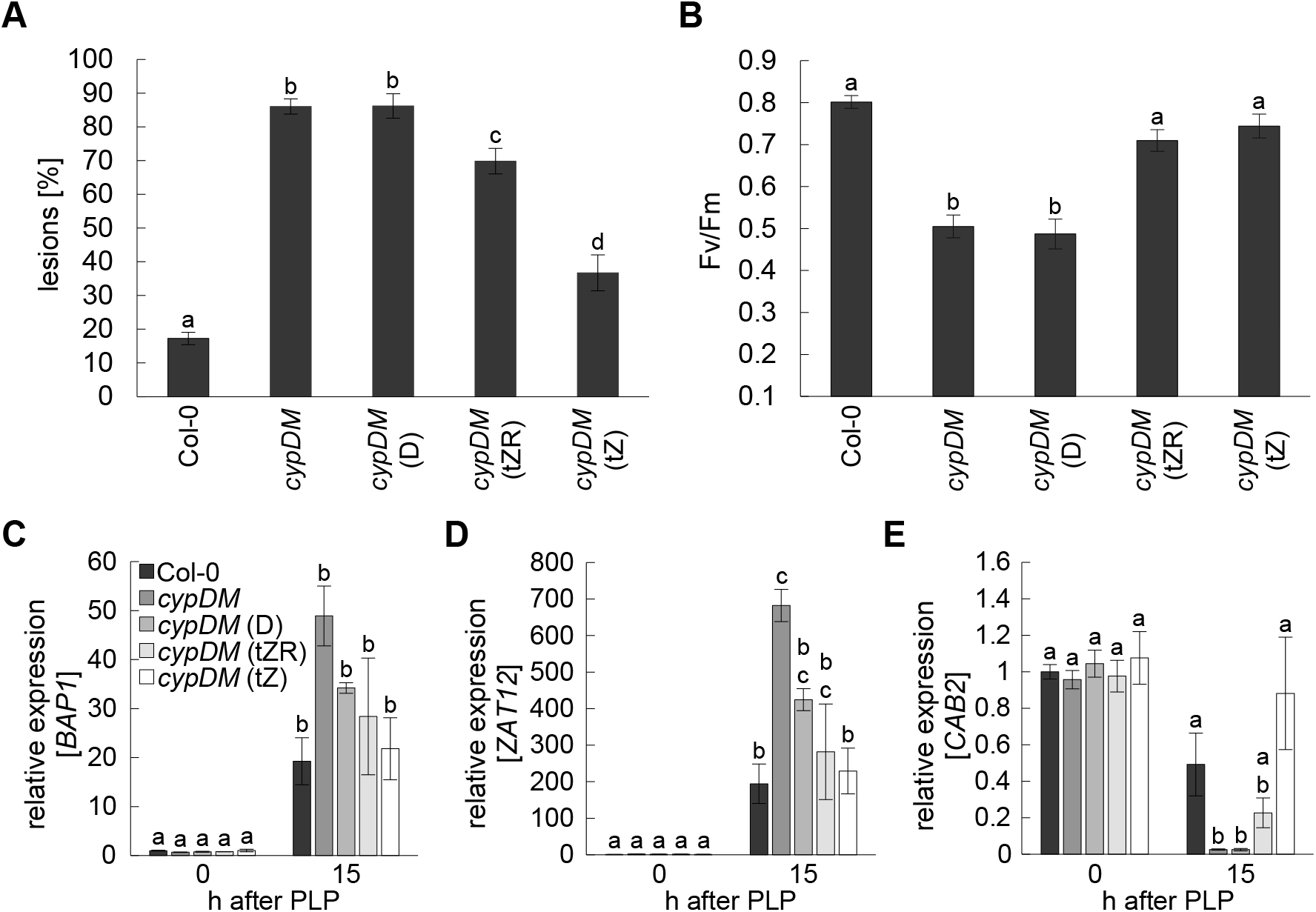
Pretreatment of CK-deficient plants with *t*Z-type CKs reduces the damage caused by photoperiod stress. *cypDM* mutant plants were watered-daily for five weeks with 10 μM *t*Z, 10 μM *t*ZR or DMSO solvent control. Thereafter, the consequences of PLP treatment on these plants were compared to untreated *cypDM* and wild-type plants. (A) Percentage of lesion formation in 5-weeks-old short day-grown plants the day after PLP treatment (one-way ANOVA; p ≤ 0.05; n = 12). (B) Photosystem II maximum quantum efficiency (F_v_/F_m_) of leaves evaluated in A (one-way ANOVA; p ≤ 0.05; n = 15). (C - E) Expression of marker genes (*BAP1, ZAT12*, *CAB2*) 0 h and 15 h after PLP treatment (one/two-way ANOVA; p ≤ 0.05; n ≥ 3). The expression level of wild type at the end of the PLP treatment (0 h) was set to 1. Abbreviations: D, DMSO; *t*Z, *trans*-zeatin; *t*ZR, *trans*-zeatin-riboside. Letters indicate statistical groups (p ≤ 0.05). Error bars indicate SE. Pictures of representative plants tested in A and B after PLP treatment are shown in Fig. S1B.

### AHP2, AHP3 and AHP5 act redundantly in photoperiod stress signaling

In *Arabidopsis*, AHK receptors transduce the CK signal to AHPs and phosphorylated AHP1 to AHP5 activate type-B ARRs (Hutchison et al., 2006). Although AHPs are involved in several developmental processes and responses to stress (Hutchison et al., 2006), their role in photoperiod stress has not been investigated so far. Thus, the *ahp2,3,5* triple mutant as well as the corresponding double mutants were exposed to photoperiod stress.

Compared to wild-type plants, about twice more leaves showed lesion formation in *ahp2,3* and *ahp2,3,5* plants (Fig. 4A, Supplemental Fig. 1C). In correspondence, the photosynthetic capacity of *ahp2,3,5* plants was decreased compared to all other genotypes (Fig. 4B). Functional redundancy of AHPs in the response to photoperiod stress was also reflected by the response of marker genes. While the stronger induction of *BAP1* and *ZAT12* expression during the night following the PLP was apparent in all *ahp* double and triple mutants compared to wild type, the amplitude was the highest in *ahp2,3* and *ahp2,3,5* (Fig. 4C, D). Similarly, a decrease of *CAB2* transcript levels (two- to three-fold) was more apparent in *ahp2,3* and *ahp2,3,5* plants than in *ahp2,5* and *ahp3,5* 15 hours after the PLP (Fig. 4E).

**Figure 4.**
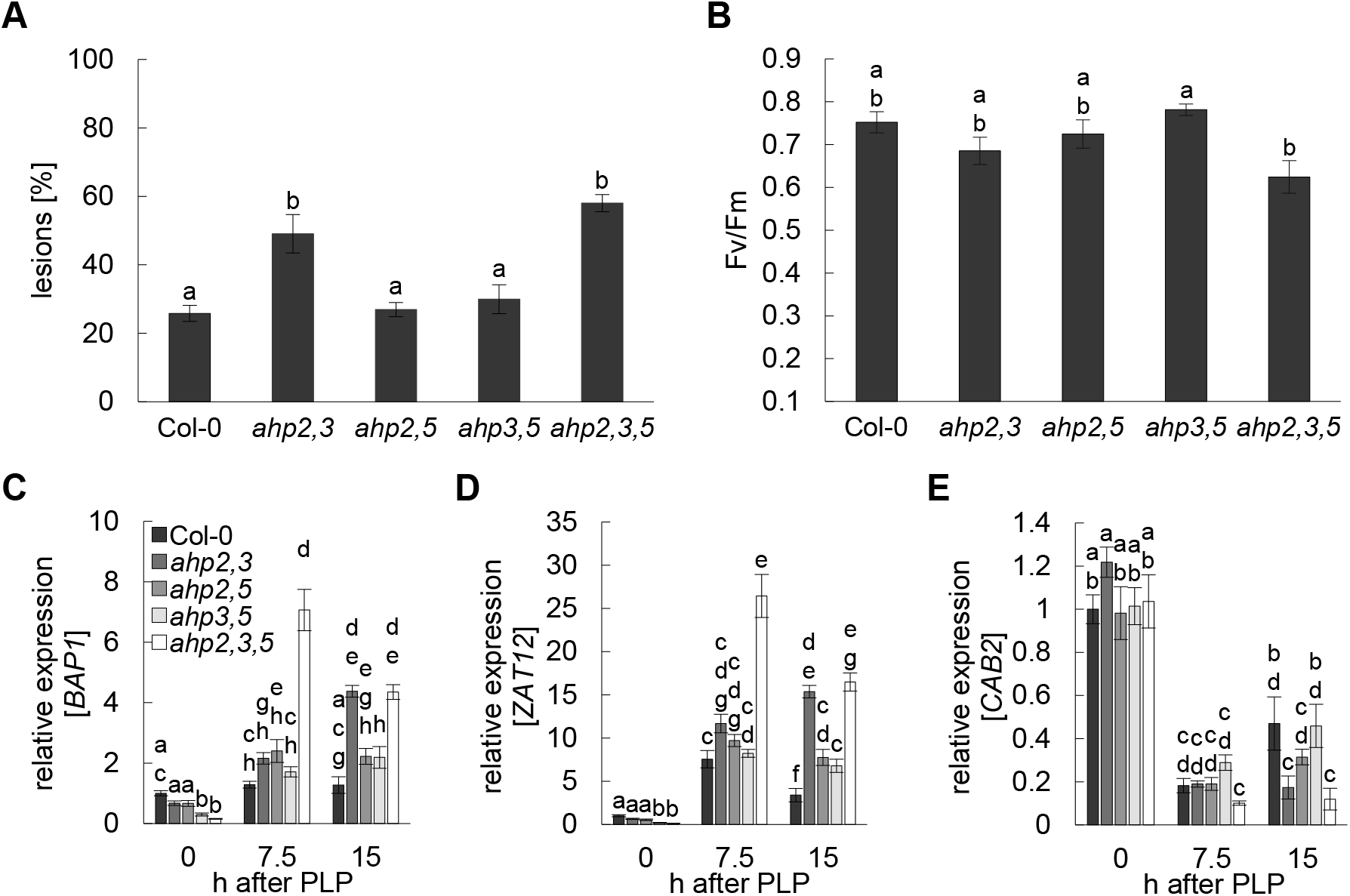
AHP2, AHP3 and AHP5 act redundantly during photoperiod stress. (A) Lesion formation in 5-weeks-old Col-0 and *ahp* mutant plants the day after PLP treatment (one-way ANOVA; p ≤ 0.05; n = 15). (B) Photosystem II maximum quantum efficiency (Fv/Fm) of leaves the day after PLP treatment (one-way ANOVA; p ≤ 0.05; n = 15). (C - E) Relative expression of marker genes (*BAP1, ZAT12*, *CAB2*) 0 h, 7.5 h and 15 h after PLP treatment. The expression level of wild type at time point 0 h was set to 1. Letters indicate statistical groups (two-way ANOVA; p ≤ 0.05; n ≥ 3). Error bars indicate SE. Pictures of representative plants tested in A and B after PLP treatment are shown in Fig. S1C.

Summing up, AHPs were shown to act redundantly in photoperiod stress signaling with AHP2 and AHP3 having a more prominent role in comparison to AHP5.

### Loss of ARR10 and ARR12 rescues the photoperiod stress sensitivity of *arr2* mutants

After phosphorylation by AHPs, type-B ARRs regulate the CK signaling output. Three members of the type-B ARR family - namely ARR2, ARR10 and ARR12 - act in photoperiod stress signaling (Nitschke et al., 2016). However, the analysis was limited to changes in Fv/Fm and the combination of all three mutant alleles was not tested. Hence, we created *arr2,10,12* triple mutant plants and exposed them to a PLP treatment along with the corresponding double and single mutants.

Consistent with the findings of Nitschke et al. (2016), the percentage of lesion forming leaves in *arr2* plants was increased 2.5-fold compared to wild-type plants after photoperiod stress treatment. In contrast, *arr10*, *arr12* and *arr10,12* mutants did not differ from wild type with respect to lesion formation (Fig. 5A, Suppl. Fig. 1D). Surprisingly, also *arr2,10* and *arr2,12* plants were indistinguishable from wild type while *arr2,10,12* plants were much more sensitive to photoperiod stress. This indicated that ARR2, ARR10 and ARR12 may interact in a complex manner to regulate the response to photoperiod stress. Measurement of the photosynthetic capacity after photoperiod stress treatment confirmed that *arr2* leaves were more affected after the PLP compared to all other genotypes except for *arr2,10,12*, which were even stronger affected (Fig. 5B). At the molecular level, the response of the different *arr* mutants varied (Fig. 5C-E). The abundance of *BAP1* and *ZAT12* did not give clear indications whether the mutants tested differed in their photoperiod stress response as the majority of differences were not statistically significant (Fig. 5C, D). In contrast, 15 h after the exposure to photoperiod stress *CAB2* was less abundant in *arr2* and *arr2,10,12* in comparison to all other genotypes (Fig. 5E). Consistent with the similar phenotypic response in terms of lesion formation and Fv/Fm, *CAB2* expression was lowered to a similar level in all other genotypes and wild type 7.5 h and 15 h after the PLP.

**Figure 5.**
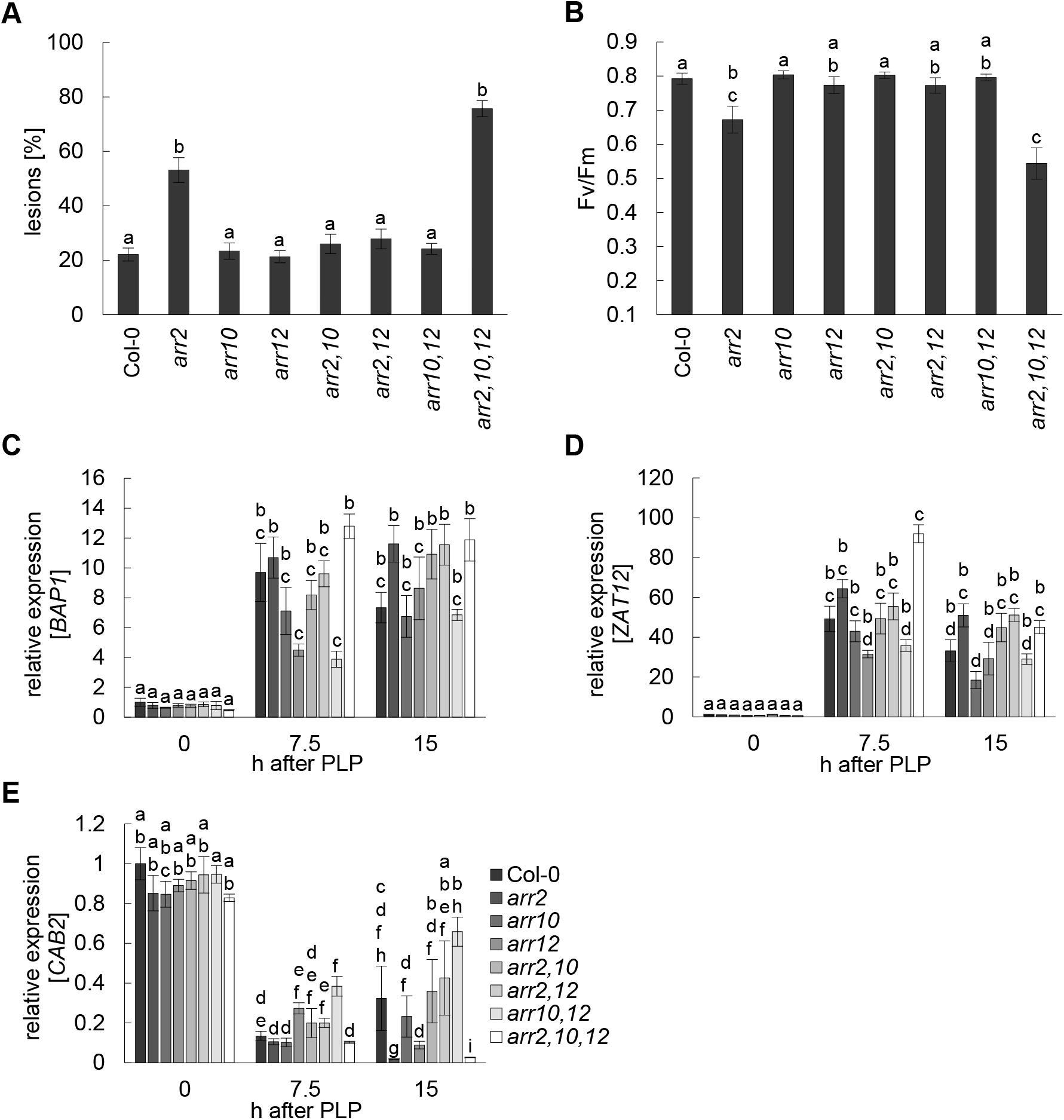
ARR2, ARR10 and ARR12 interact to respond to photoperiod stress. (A) Quantification of lesion forming leaves in 5-weeks-old Col-0 and type-B ARR mutants the day after the PLP treatment (one-way ANOVA; p ≤ 0.05; n = 15). (B) Photosystem II maximum quantum efficiency (Fv/Fm) of leaves the day after PLP treatment (one-way ANOVA; p ≤ 0.05; n = 15). (C - E) Relative expression of marker genes (*BAP1, ZAT12*, *CAB2*) 0 h, 7.5 h and 15 h after PLP treatment. The expression level of wild type at the end of the PLP treatment (0 h) was set to 1. Letters indicate statistical groups (two-way ANOVA/Paired Wilcoxon test; p ≤ 0.05; n ≥ 3). Error bars indicate SE. Pictures of representative plants tested in A and B after PLP treatment are depicted in Fig. S1D.

In summary, the results confirmed the results of Nitschke et al. (2016) who reported a positive regulatory function of ARR2 in photoperiod stress. In addition, the results suggested that ARR2, ARR10 and ARR12 interact in a complex manner to regulate the response to photoperiod stress.

## DISCUSSION

### The CK concentration is increased in response to photoperiod stress

Here we reported on the functional relevance of root-derived CK in the response to photoperiod stress. Wild-type plants grown under short-day conditions and experiencing a PLP responded by increasing the CK concentration in their leaves (Fig.1, Table S1). As plants with a reduced CK concentration or signaling are particularly sensitive to photoperiod stress (Nitschke et al., 2016), this response may be part of a defense mechanism enabling wild-type plants to react appropriately to photoperiod stress and to cope with its consequences. Altered CK concentrations are often part of the response to abiotic stress and they may either increase or decrease. A decreased CK concentration was found after exposure to several abiotic stresses like heat, salt or drought stress (Bano, Hansen, Dörffling, & Hahn, 1994; Caers, Rüdelsheim, van Onckelen, & Horemans, 1985; Itai, Ben-Zioni, & Ordin, 1973; Nishiyama et al., 2011). Plants with a lower CK status were more stress resistant indicating a functional relevance of the reduced CK level (Nishiyama et al., 2011). In contrast, under high light stress CK has a protective function. It represses excessive starch grain and plastoglobuli formation and is required for a functional D1 repair cycle (Cortleven et al., 2014). In the response to biotic stress such as *Pseudomonas* infection, CK is required for an effective defense regulating the oxidative burst through ARR2 (Arnaud et al., 2017; Choi et al., 2010). CK is known also from other instances to regulate the response to oxidative stress (Pavlů et al., 2018; Cortleven et al., 2019) which is a hallmark of the response to photoperiod stress (Nitschke et al., 2016). We propose that one function of the enhanced CK formation could be to properly respond to oxidative stress caused by PLP treatment (Nitschke et al., 2016).

### Root-derived *t*Z-type CKs act as protectants against photoperiod stress

Root-derived *t*Z-type CKs were shown to be the most relevant CK type for the response to photoperiod stress (Fig. 2). The major transport form, *t*ZR, as well as to a minor extent its bioactive derivative *t*Z, are transported from the root to the shoot via the xylem flow requiring the transporter ABCG14 (Ko et al., 2014; Zhang et al., 2014). *abcg14* mutants are thus deficient in *tZ* in the shoot (Ko et al., 2014; Zhang et al., 2014) and consistently these mutants showed a very strong response to photoperiod stress. In *cypDM* mutants, the lower levels of *t*Z-type CKs in the shoot are compensated by an increased level of iP-type CK (Kiba et al., 2013). The inability of these higher levels of iP-type CKs to compensate the sensitive photoperiod stress response of *cypDM* mutants corroborates the functional relevance of *t*Z-type CKs. Consistent with a major role of *t*Z-type CKs is also the functional relevance of AHK3 in photoperiod stress signaling (Nitschke et al., 2016). AHK3 displays an about tenfold higher sensitivity to *t*Z than to iP while AHK2 and AHK4/CRE1 have similar affinities to both iP and *t*Z (Lomin et al., 2015; Romanov, Lomin, & Schmülling, 2006; Stolz et al., 2011). It has been proposed that the affinity profile of AHK3 is particularly set to respond to root-derived CK (Romanov et al., 2006).

Further support for a role of root-derived CK in photoperiod stress protection came from supplementation experiments. Watering of *cypDM* plants with either *t*ZR or *t*Z demonstrated that both metabolites can protect plants against photoperiod stress although *t*Z was more effective (Fig. 3). Both *t*Z and *t*ZR supplementation rescued the decrease in type-A *ARR* transcript abundance in these plants demonstrating that after application through roots they reached the shoot in a biologically effective concentration. Different roles for root-derived *t*Z and *t*ZR have been reported by Osugi et al. (2017). It might be that the ability of certain tissues to convert inactive *t*ZR to active *t*Z, as discussed in Romanov et al. (2018), might have an impact on the plant’s response to photoperiod stress.

The functional relevance of root-derived CK in the response to photoperiod stress raises the question how information about a stress perceived and acting primarily in the shoot is relayed to the root. One possibility is that the light signal is perceived and interpreted in the root directly (Sun, Yoda, & Suzuki, 2005; Sun, Yoda, Suzuki, & Suzuki, 2003). Another possibility is that an instructive chemical signal is formed in the shoot and transported to the root to induce synthesis of *t*Z CK. This signal could be iP-type CK as these are mainly formed in the shoot and known to be transported to the root through the phloem (Hirose et al., 2008; Kudo, Kiba, & Sakakibara, 2010). iP-type CKs could then positively regulate *t*Z-type CK formation as they not only serve as a precursor for *t*Z-formation but also induce the expression of *CYP735A2* (Takei et al., 2004). Another candidate for a chemical signal is jasmonic acid which is increased in response to photoperiod stress in sensitive genotypes (Nitschke et al., 2016) and which has recently been shown to be a shoot-to-root signal (Schulze et al., 2019).

### ARR2, ARR10 and ARR12 regulate the response to photoperiod stress in a complex manner

AHP2, AHP3 and AHP5 act redundantly in the response to photoperiod stress (Fig.4). The functional redundancy of these AHPs has been shown before in the context of seed, primary root and hypocotyl development (Hutchison et al., 2006). Our results integrate AHPs into the CK-dependent photoperiod stress signaling pathway that so far involved AHK3 and ARR2 as the main signaling components (Nitschke et al., 2016).

Downstream of the AHPs act several transcription factors to realize the transcriptional output of the photoperiod stress response. ARR2 has a predominant role in mediating CK activity in leaves (Hwang & Sheen, 2001) but its redundant function with ARR10 and ARR12 has not yet been described. The latter two ARRs are better known for their role in regulating most CK-related vegetative developmental processes together with ARR1 (Argyros et al., 2008; Ishida, Yamashino, Yokoyama, & Mizuno, 2008). Analysis of single and double mutants showed that loss of either *ARR10* or *ARR12* rescued the stress phenotype of *arr2* plants while the loss of both factors enhanced the stress response of *arr2* (Fig. 5). This hints to a complex regulatory mechanism between these three transcription factors during photoperiod stress signaling. A complex relationship among these type-B ARRs has also been described for their role in regulating root elongation. *arr12* and *arr10,12* root elongation was less affected by CK treatment than that of *arr2,12* and *arr2,10,12* (Mason et al., 2005). For type-B ARR dependent gene regulation, a model has been proposed in which simultaneous binding of multiple/different type-B ARRs and unknown factors to certain promoter regions is crucial (Ramireddy, Brenner, Pfeifer, Heyl, & Schmülling, 2013). However, experimental evidence for a direct interaction between members of the type-B ARR family is rare. An interaction of ARR2 and ARR14 has been described using a two-hybrid system in yeast (Dortay, Mehnert, Bürkle, Schmülling, & Heyl, 2006). Recently, it was found that the C-termini of ARR1 and ARR12 interact to regulate auxin synthesis (Yan et al., 2017). It could also be that interactions between type-B ARRs are context-dependent as it is known for the phosphorylation-dependent homodimerization of bacterial RRs (Mack, Gao, & Stock, 2009). Similarly, ARR18 can homodimerize when both ARR18 proteins are either both phosphorylated or both not phosphorylated (Veerabagu et al., 2012).

The different phenotypic and in part molecular responses to photoperiod stress of *arr* mutants could be explained by a model, in which ARR2, ARR10 and ARR12 interact with a yet unknown interaction partner (X) that is essential for photoperiod stress resistance (Fig. 6). It is predicted that the affinity of ARR2 to X would be higher than the affinities of ARR10 and ARR12 to X. In addition, we propose a direct or indirect interaction of ARR10 and ARR12. In photoperiod stress-treated wild-type plants, ARR2 would interact with X resulting in photoperiod stress resistance while ARR10 and ARR12 together would have independent auxiliary functions. In *arr2* plants, X would not have an interaction partner and thus would be unable to function in stress protection because ARR10 and ARR12 would not be available as interaction partners. Consequently, stress resistance would be lowered. Resistance of *arr2,10* and *arr2,12* plants would be caused by the loss of the ARR10-ARR12 association and the resulting interaction of X with ARR10 or ARR12. Ultimately, the enhanced stress phenotype of *arr2,10,12* plants would be caused by the complete loss of interaction partners for X.

**Figure 6.**
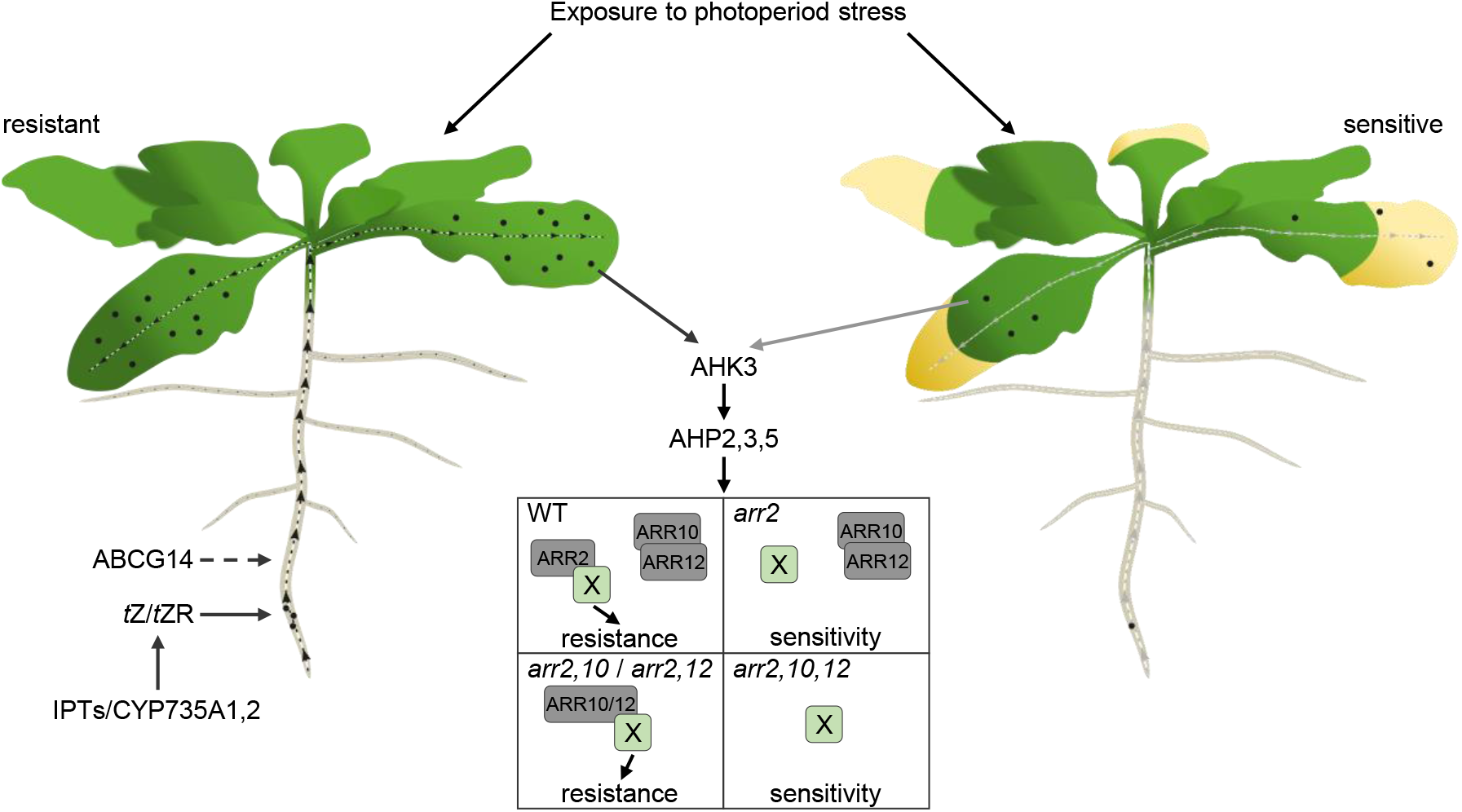
Model showing the role of CK in regulating the response to photoperiod stress. During exposure to photoperiod stress, wild-type plants (left) increase their CK levels. IPT and CYP735A proteins increase synthesis of *t*Z-type CK (black balls) in roots which are transported via ABCG14 to the shoot (black dashed line) where they activate CK signaling mainly through AHK3. AHP2, AHP3 and AHP5, and ARR2, ARR10 and ARR12. Impairment of either *t*Z-type CK synthesis or transport (less molecules and grey dashed lines) induce weaker CK signaling causing higher sensitivity to photoperiod stress (right plant). The central four rectangles show a model for type-B ARR-dependent regulation of the response. It is proposed that ARR2, ARR10 and ARR12 interact in the wild type (WT) with a yet unknown interaction partner (X) essential for photoperiod stress resistance (rectangle top left). The affinity of ARR2 to X is higher than the affinities of ARR10 and ARR12 to X. Additionally, ARR10 and ARR12 directly or indirectly interact with each other. In *arr2* plants (rectangle top right), X does not have an interaction partner and thus would be unable to function while ARR10 and ARR12 still interact with each other leading to the formation of the photoperiod stress syndrome. Resistance of *arr2,10* and *arr2,12* plants (rectangle bottom left) is caused by the loss of ARR10-ARR12 association and the resulting interaction of X with ARR10 or ARR12. Ultimately, the enhanced photoperiod stress sensitivity of *arr2,10,12* plants (rectangle bottom right) would be caused by the complete loss of interaction partners for X.

Beside the interaction amongst ARRs, interactions between several type-B ARRs and other proteins exist. For example, ARR1, ARR2 and ARR14 interact with the DELLA proteins RGA1 and GAI to regulate root development and photomorphogenesis (Marín-de la Rosa et al., 2015; Yan et al., 2017). During the regulation of auxin synthesis, EIN3 interacts with the C-terminus of ARR1 and thereby increases ARR1 activity (Yan et al., 2017). As part of the crosstalk between CK and abscisic acid, ARR1, ARR11 and ARR12 directly interact with SUCROSE NON-FERMENTING-1 (SNF1)- RELATED PROTEIN KINASE2 (SnRK2) kinases and thereby inhibit their function prior to drought stress (Huang et al., 2018). Future experiments might resolve whether and how type-B ARRs interact with each other or with other proteins during photoperiod stress.

## Supporting information

Frank et al_Supplemental data

## SUPPORTING INFORMATION

**Table S1.** Changes in CK concentration by PLP treatment. The indicated time points (control/PLP 1 to 5) correspond to those shown in Figure 1A. Bold numbers indicate statistically significant difference in PLP samples compared to the respective controls at the same time point in a paired Student’s *t*-test (p ≤ 0.05). Values are given as pmol g^−1^ FW ± SD (n = 5). Concentrations below detection limit are referred to as <LOD. RMP, riboside monophosphates; *O*G, *O*-glucosides; R*O*G, riboside-*O*-glucoside; 7G, 7-glucoside; 9G, 9-glucoside.

**Figure S1.** Representative plants after PLP treatment. The pictures illustrate the phenotype of plants used for experiments shown in Fig. 2 to Fig. 5. Pictures were taken two days after PLP treatment and belong to Fig. 2 (A), Fig. 3 (B), Fig. 4 (C), and Fig. 5 (D). Details of the experiments can be found in the legends of the respective figures.

**Figure S2.** Pretreatment of CK-deficient plants with *t*Z-type CKs rescues differential expression of CK response genes. Expression of *ARR5* (A) and *ARR6* (B) 0 h after PLP treatment relative to wild type. Letters indicate statistical groups (one-way ANOVA; p ≤ 0.05; p ≤ 0.05; n ≥ 3).

## ACKNOWLEDGEMENTS

We thank Sören Werner and Gabi Grüschow for generating the *arr* mutants and Hana Martínková and Petra Amakorová for technical assistance with cytokinin profiling. We acknowledge funding by the Deutsche Forschungsgemeinschaft (DFG) (grant Schm 814-27/1 and Collaborative Research Centre 973, www.sfb.973). This work was supported by the Czech Science Foundation (No. 19-00973S) and from the European Regional Development Fund-Project “Plants as a tool for sustainable global development” (No. CZ.02.1.01/0.0/0.0/16_019/0000827).

## AUTHOR CONTRIBUTIONS

MF, AC and TS developed the project; MF designed and performed experiments, partly together with AC; MF, AC and TS analyzed data; ON measured cytokinin concentrations, MF, AC and TS wrote the article, all other authors read and contributed to previous versions and approved the final version.

