## Supplementary material for "Root-derived *trans*-zeatin cytokinin protects *Arabidopsis* plants against photoperiod stress": Frank et al_Supplemental data

**Table S1.** Changes in CK concentration by PLP treatment. The indicated time points (control/PLP 1 to 5) correspond to those shown in Figure 1A. Bold numbers indicate statistically significant difference in PLP samples compared to the respective controls at the same time point in a paired Student's t-test ($p \leq 0.05$). Values are given as pmol g-1 FW ± SD (n = 5). Concentrations below detection limit are referred to as <LOD. RMP, riboside monophosphates; *O*G, *O*-glucosides; R*O*G, riboside-*O*-glucoside; 7G, 7-glucoside; 9G, 9-glucoside.

| | **1** | | **2** | | **3** | | **4** | | **5** | |
|---|---|---|---|---|---|---|---|---|---|---|
| | **control** | **PLP** | **control** | **PLP** | **control** | **PLP** | **control** | **PLP** | **control** | **PLP** |
| **iP** | 0.090±0.027 | 0.103±0.021 | 0.078±0.024 | **0.161±0.034** | 0.115±0.018 | **0.193±0.056** | 0.074±0.022 | **0.234±0.059** | 0.063±0.014 | 0.059±0.019 |
| **iPR** | 0.24 ± 0.06 | **0.75 ± 0.11** | 0.40 ± 0.10 | **0.87 ± 0.10** | 0.51 ± 0.05 | 0.61 ± 0.13 | 0.27 ± 0.05 | **1.11 ± 0.25** | 0.56 ± 0.16 | 0.54 ± 0.17 |
| **iPRMP** | 5.13 ± 0.74 | **21.61± 1.81** | 10.86± 1.77 | **20.66± 1.46** | 10.98± 1.02 | 14.69± 3.15 | 5.27 ± 1.22 | 6.56 ± 1.97 | 14.44± 1.19 | **10.32± 2.84** |
| **iP7G** | 22.04± 1.25 | 23.56± 2.89 | 20.03± 0.74 | 21.10± 1.31 | 23.19± 1.66 | 23.91± 0.85 | 20.64± 0.58 | **24.65± 1.92** | 21.19± 0.98 | 20.59± 2.74 |
| **iP9G** | 1.90 ± 0.11 | 2.10 ± 0.25 | 1.70 ± 0.04 | 1.91 ± 0.19 | 2.01 ± 0.22 | 2.04 ± 0.07 | 1.78 ± 0.11 | 1.96 ± 0.18 | 1.76 ± 0.10 | 1.58 ± 0.23 |
| ***t*Z** | 0.009±0.003 | 0.009±0.002 | 0.007±0.001 | **0.010±0.002** | 0.003±0.001 | **0.006±0.001** | 0.006±0.001 | 0.007±0.001 | 0.004±0.001 | 0.003±0.001 |
| ***t*ZR** | 1.67 ± 0.19 | **3.03 ± 0.53** | 2.43 ± 0.43 | **3.51 ± 0.77** | 2.90 ± 0.90 | 4.04 ± 0.89 | 1.90 ± 0.24 | **3.36 ± 0.78** | 2.59 ± 0.47 | 2.73 ± 0.46 |
| ***t*ZRMP** | 8.11 ± 1.20 | **13.25± 3.13** | 7.56 ± 1.65 | **11.89± 2.17** | 6.01 ± 0.31 | **8.88 ± 1.75** | 6.04 ± 1.13 | 4.83 ± 1.25 | 7.85 ± 1.61 | **3.98 ± 0.91** |
| ***t*Z*O*G** | 6.77 ± 0.43 | 6.47 ± 0.49 | 5.64 ± 0.31 | 6.02 ± 0.24 | 5.37 ± 0.17 | 5.88 ± 0.26 | 5.52 ± 0.46 | **6.42 ± 0.61** | 5.52 ± 0.16 | **4.67 ± 0.57** |
| ***t*ZR*O*G** | 1.10 ± 0.05 | 1.08 ± 0.11 | 0.98 ± 0.08 | 1.06 ± 0.05 | 0.90 ± 0.04 | **1.24 ± 0.09** | 1.05 ± 0.08 | **1.46 ± 0.17** | 0.93 ± 0.04 | 0.90 ± 0.15 |
| ***t*Z7G** | 87.53± 3.94 | 81.79± 6.96 | 74.79± 5.97 | 79.45± 2.35 | 77.34± 2.45 | 76.07± 2.10 | 79.79± 6.84 | 85.25± 8.16 | 73.74± 1.31 | **63.65± 6.76** |
| ***t*Z9G** | 26.97± 0.60 | **23.71± 2.26** | 22.16± 1.99 | 22.78± 0.76 | 20.82± 1.05 | 20.23± 0.73 | 22.18± 2.32 | 24.80± 2.21 | 19.62± 1.60 | 16.82± 2.49 |
| **DHZ** | <LOD | <LOD | <LOD | <LOD | <LOD | <LOD | <LOD | <LOD | <LOD | <LOD |
| **DHZR** | 0.026±0.007 | **0.053±0.015** | 0.027±0.008 | **0.049±0.010** | 0.031±0.009 | **0.075±0.021** | 0.023±0.002 | **0.138±0.037** | 0.023±0.005 | **0.067±0.020** |
| **DHZRMP** | <LOD | <LOD | <LOD | <LOD | <LOD | <LOD | <LOD | <LOD | <LOD | <LOD |
| **DHZ*O*G** | 0.059±0.005 | 0.057±0.003 | 0.054±0.002 | 0.055±0.006 | 0.048±0.006 | **0.062±0.005** | 0.042±0.006 | **0.064±0.006** | 0.041±0.003 | 0.051±0.008 |
| **DHZR*O*G** | 0.059±0.005 | 0.062±0.013 | 0.067±0.006 | 0.072±0.015 | 0.057±0.009 | **0.096±0.021** | 0.056±0.007 | **0.122±0.036** | 0.056±0.010 | 0.068±0.017 |
| **DHZ7G** | 5.73 ± 0.17 | 5.56 ± 0.43 | 5.13 ± 0.20 | 5.58 ± 0.39 | 5.11 ± 0.20 | **6.11 ± 0.35** | 4.70 ± 0.28 | **6.02 ± 0.86** | 4.46 ± 0.24 | 4.38 ± 0.54 |
| **DHZ9G** | 0.17 ± 0.03 | 0.14 ± 0.01 | 0.14 ± 0.01 | 0.12 ± 0.01 | 0.11 ± 0.02 | 0.10 ± 0.01 | 0.11 ± 0.01 | **0.16 ± 0.04** | 0.11 ± 0.01 | 0.10 ± 0.02 |
| ***c*Z** | <LOD | <LOD | <LOD | <LOD | <LOD | <LOD | <LOD | <LOD | <LOD | <LOD |
| ***c*ZR** | 0.41 ± 0.04 | **0.10 ± 0.02** | 0.30 ± 0.10 | 0.21 ± 0.06 | 0.33 ± 0.02 | 0.34 ± 0.03 | 0.44 ± 0.01 | **3.22 ± 0.92** | 0.16 ± 0.03 | **1.55 ± 0.46** |
| ***c*ZRMP** | 3.72 ± 0.58 | **1.54 ± 0.15** | 2.84 ± 0.42 | 2.22 ± 0.46 | 4.39 ± 0.23 | 4.43 ± 0.57 | 3.56 ± 0.19 | **6.18 ± 0.81** | 1.74 ± 0.34 | **4.62 ± 0.88** |
| ***c*Z*O*G** | 1.14 ± 0.08 | 1.20 ± 0.12 | 1.02 ± 0.07 | **1.18 ± 0.09** | 1.16 ± 0.11 | 1.29 ± 0.06 | 0.99 ± 0.04 | **1.29 ± 0.04** | 1.04 ± 0.08 | **1.72 ± 0.42** |
| ***c*ZR*O*G** | 2.73 ± 0.16 | 2.48 ± 0.21 | 2.30 ± 0.13 | 2.07 ± 0.17 | 2.64 ± 0.12 | **3.20 ± 0.30** | 2.73 ± 0.11 | 3.17 ± 0.38 | 2.54 ± 0.25 | 2.56 ± 0.31 |
| ***c*Z7G** | 13.32± 0.99 | 12.53± 1.36 | 10.46± 0.46 | 10.82± 1.04 | 12.88± 0.79 | 12.59± 0.93 | 12.13± 1.30 | 10.67± 1.22 | 11.46± 1.05 | **9.83 ± 0.73** |
| ***c*Z9G** | 0.28 ± 0.01 | **0.23 ± 0.03** | 0.21 ± 0.02 | **0.17 ± 0.02** | 0.24 ± 0.03 | **0.17 ± 0.01** | 0.22 ± 0.02 | 0.20 ± 0.02 | 0.25 ± 0.02 | **0.18 ± 0.02** |

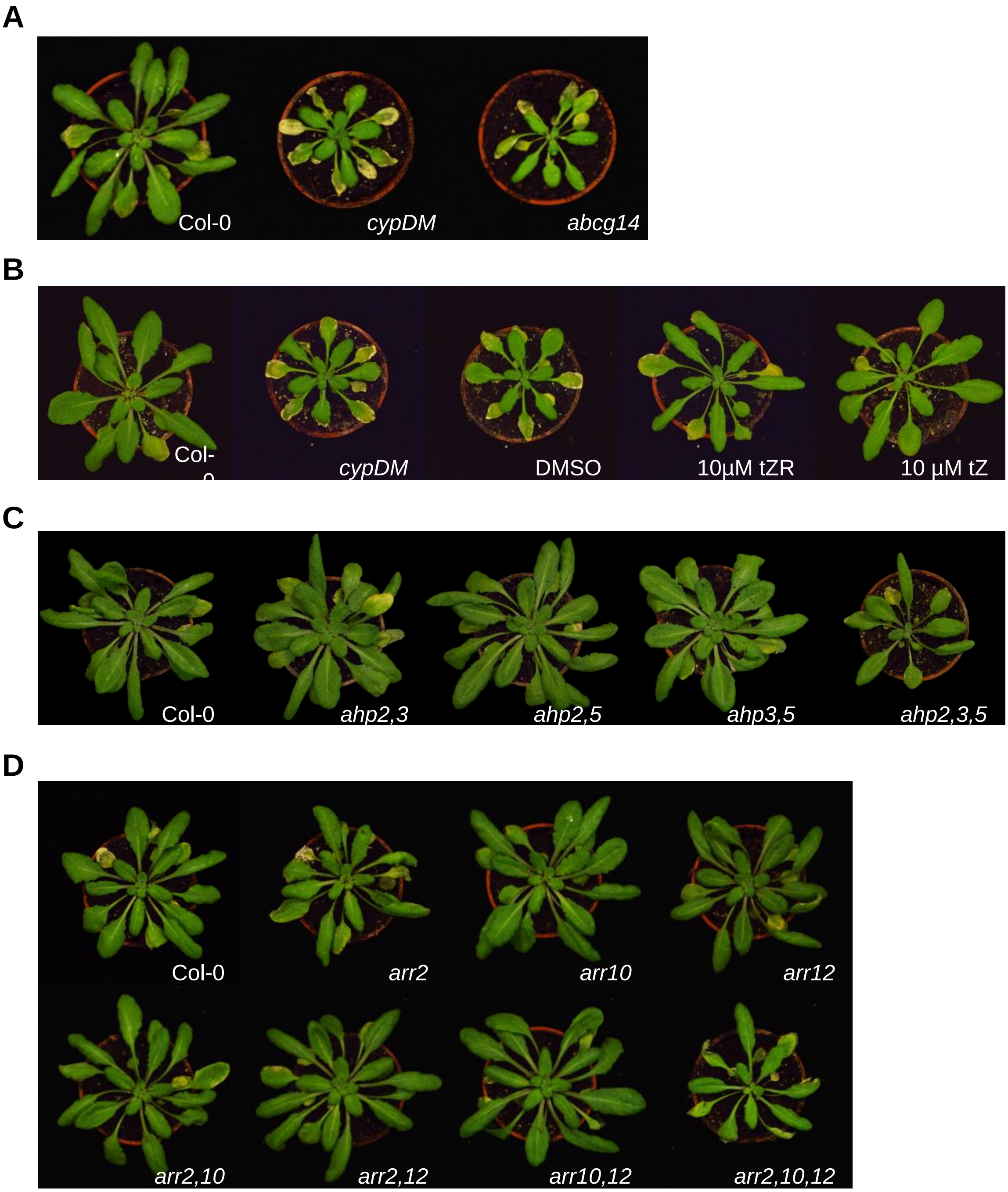


**Figure S1.** Representative plants after PLP treatment. The pictures illustrate the phenotype of plants used for experiments shown in Fig. 2 to Fig. 5. Pictures were taken two days after PLP treatment and belong to Fig. 2 (A), Fig. 3 (B), Fig. 4 (C), and Fig. 5 (D). Details of the experiments can be found in the legends of the respective figures.

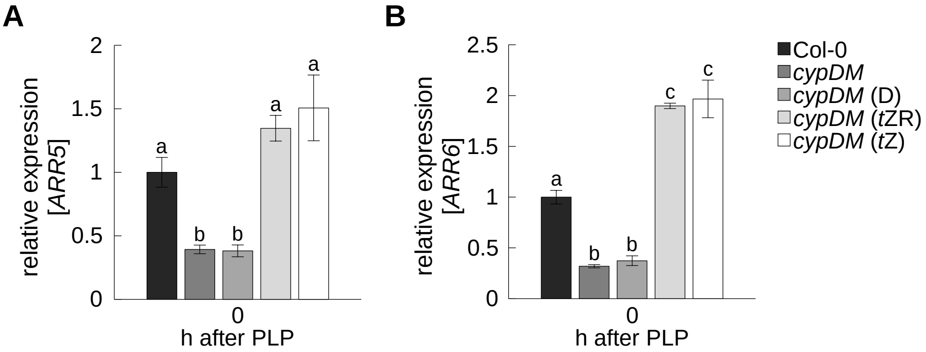


**Figure S2.** Pretreatment of CK-deficient plants with *t*Z-type CKs rescues differential expression of CK response genes. Expression of ARR5 (A) and ARR6 (B) 0 h after PLP treatment relative to wild type . Letters indicate statistical groups (one-way ANOVA; $p \leq 0.05$; $p \leq 0.05$; $n \geq 3$).
